# Environmental responses, not species interactions, determine synchrony of dominant species in semiarid grasslands

**DOI:** 10.1101/047480

**Authors:** Andrew T. Tredennick, Claire de Mazancourt, Michel Loreau, Peter B. Adler

## Abstract

Temporal asynchrony among species helps diversity to stabilize ecosystem functioning, but identifying the mechanisms that determine synchrony remains a challenge. Here, we refine and test theory showing that synchrony depends on three factors: species responses to environmental variation, interspecific interactions, and demographic stochasticity. We then conduct simulation experiments with empirical population models to quantify the relative influence of these factors on the synchrony of dominant species in five semiarid grasslands. We found that the average synchrony of per capita growth rates, which can range from 0 (perfect asynchrony) to 1 (perfect synchrony), was higher when environmental variation was present (0.62) rather than absent (0.43). Removing interspecific interactions and demographic stochasticity had small effects on synchrony. For the dominant species in these plant communities, where species interactions and demographic stochasticity have little influence, synchrony reflects the covariance in species responses to the environment.

## INTRODUCTION

Ecosystems are being transformed by species extinctions (Cardinale et al. 2012), changes in community composition (Vellend et al. 2013, Dornelas et al. 2014), and anthropogenic environmental change (Vitousek et al. 1997), impacting the provisioning and stability of ecosystem services (Loreau et al. 2001, Hooper et al. 2005, Rockstrom et al. 2009). Experiments have provided compelling evidence that decreases in species richness will decrease productivity (Tilman et al. 2001) and the temporal stability of productivity (Tilman et al. 2006, Hector et al. 2010). The stabilizing effect of species richness arises from a combination of selection effects and complementarity (Loreau and Hector 2001). Selection effects occur when a dominant species has lower than average temporal variability, which generates a positive effect on ecosystem stability (e.g., Grman et al. 2010). Complementarity occurs when species have unique temporal dynamics, causing their abundances to fluctuate asynchronously, increasing ecosystem stability. The premise of this paper is that understanding the mechanisms driving species’ temporal dynamics, and resulting (a)synchrony, is necessary to predict the impacts of global change on ecosystem stability.

Asynchronous dynamics, also known as compensatory dynamics (Gonzalez and Loreau 2009), occur whenever species synchrony is not perfect and result from individual species responding in different ways to environmental fluctuations, random chance events, and/or competitive interactions (Isbell et al. 2009, Hector et al. 2010, de Mazancourt et al. 2013, Gross et al. 2014). Species richness affects the degree of synchrony in a community because larger species pools are more likely to contain species that respond dissimilarly to environmental conditions, reducing synchrony and increasing stability (Yachi and Loreau 1999). Species richness can also affect synchrony if the strength of species interactions varies systematically with richness, because competition generally increases synchrony and reduces stability (Loreau and de Mazancourt 2013, but see Gross et al. 2014).

The effects of environmental change and species losses on ecosystem stability will depend on whether synchrony is driven by species-specific responses to environmental conditions or interspecific competition (Hautier et al. 2014). If responses to environment are important, then environmental change could alter synchrony and stability. If competition is important, then the direct effects of environmental change may not affect synchrony and stability, but species gains/losses will.

The relative role of environmental responses and competition in driving synchrony in natural plant communities remains controversial (reviewed in Gonzalez and Loreau 2009). One source of the controversy is that quantifying the relative strengths of each driver based on the covariance matrix of species abundances (e.g., Houlahan et al. 2007) is impossible. This is because observed synchrony can arise from non-unique combinations of factors (Ranta et al. 2008). For example, weak synchrony of population abundances could reflect positive environmental correlations (synchronizing effect) offset by strong competition (desynchronizing effect), or negative environmental correlations and weak competition.

Theory can help us resolve this empirical question. Recent theoretical work has identified three determinants of species synchrony: environmental stochasticity, interspecific interactions, and demographic stochasticity (Loreau and de Mazancourt 2008, 2013, Gonzalez and Loreau 2009). This theory has been developed by focusing on simple limiting cases in which only one of these three drivers operates. For example, in a community composed of large populations (no demographic stochasticity) with weak interspecific interations, community-wide species synchrony should be determined by the covariance of species’ responses to the environment (Loreau and de Mazancourt 2008). However, this prediction relies on a relatively simple population model and depends on two restrictive assumptions: (i) species’ responses to the environment are similar in magnitude and (ii) all species have similar growth rates. Whether such theoretical predictions hold in natural communities where species differences are unlikely to be symmetrical is unkown because few studies have explicitly tested theory on the drivers of species synchrony in natural communities (Mutshinda et al. 2009, Thibaut et al. 2012), and they did not consider demographic stochasiticity.

An intuitive way to quantify the effects of environmental stochasticity, demographic stochasticity, and interspecific interactions is to remove them one-by-one, and in combination. If synchrony changes more when we remove environmental stochasticity than when we remove interspecific interactions, we would conclude that environmental fluctuations are the more important driver. In principle, this could be done in an extremely controlled laboratory setting (e.g., Venail et al. 2013), but empirically-based models of interacting populations, fit with data sets from natural communities, offer a practical alternative. For example, Mutshinda et al. (2009) fit a dynamic population model to several community time series of insect and bird abundances. They used a statistical technique to decompose temporal variation into competition and environmental components, and found that positively correlated environmental responses among species determined community dynamics. Thibaut et al. (2012) used a similar approach for reef fish and came to a similar conclusion: environmental responses determine synchrony.

While a major step forward, Mutshinda et al.’s (2009) and Thibaut et al.’s (2012) modeling technique does have some limitations. First, although both studies quantified the relative importance of environmental stochasticity and interspecific interactions to explain the observed variation of species synchrony, they did not use the model to quantify how much synchrony would change if each factor were removed. Second, they relied on popluation abundance data that may or may not reliably capture competitive interactions occuring at the individual level. Third, fluctuations in abundance may mask the mechanisms that underpin species synchrony. The synchrony of species’ abundances ultimately determines the stability of total community biomass, but the processes that drive species synchrony are most tightly linked to each species’ immediate response to environmental conditions and competition (Loreau and de Mazancourt 2008). Therefore, we focus on per capita growth rates, which represent a species’ immediate response to interannual fluctuations.

Here, we use multispecies population models fit to long-term demographic data from five semi-arid plant communities to test theory on the drivers of species synchrony (Fig. 1). Our objectives are to (1) derive and test theoretical predictions of species synchrony and (2) determine the relative influence of environmental stochasticity, interspecific interactions, and demographic stochasticity on the synchrony of dominant species in natural plant communities. While our focus is limited to dominant species due to data constraints, previous work indicates that the dynamics of dominant species determine ecosystem functioning in grasslands (Smith and Knapp 2003, Bai et al. 2004, Grman et al. 2010, Sasaki and Lauenroth 2011).

**Figure 1:**
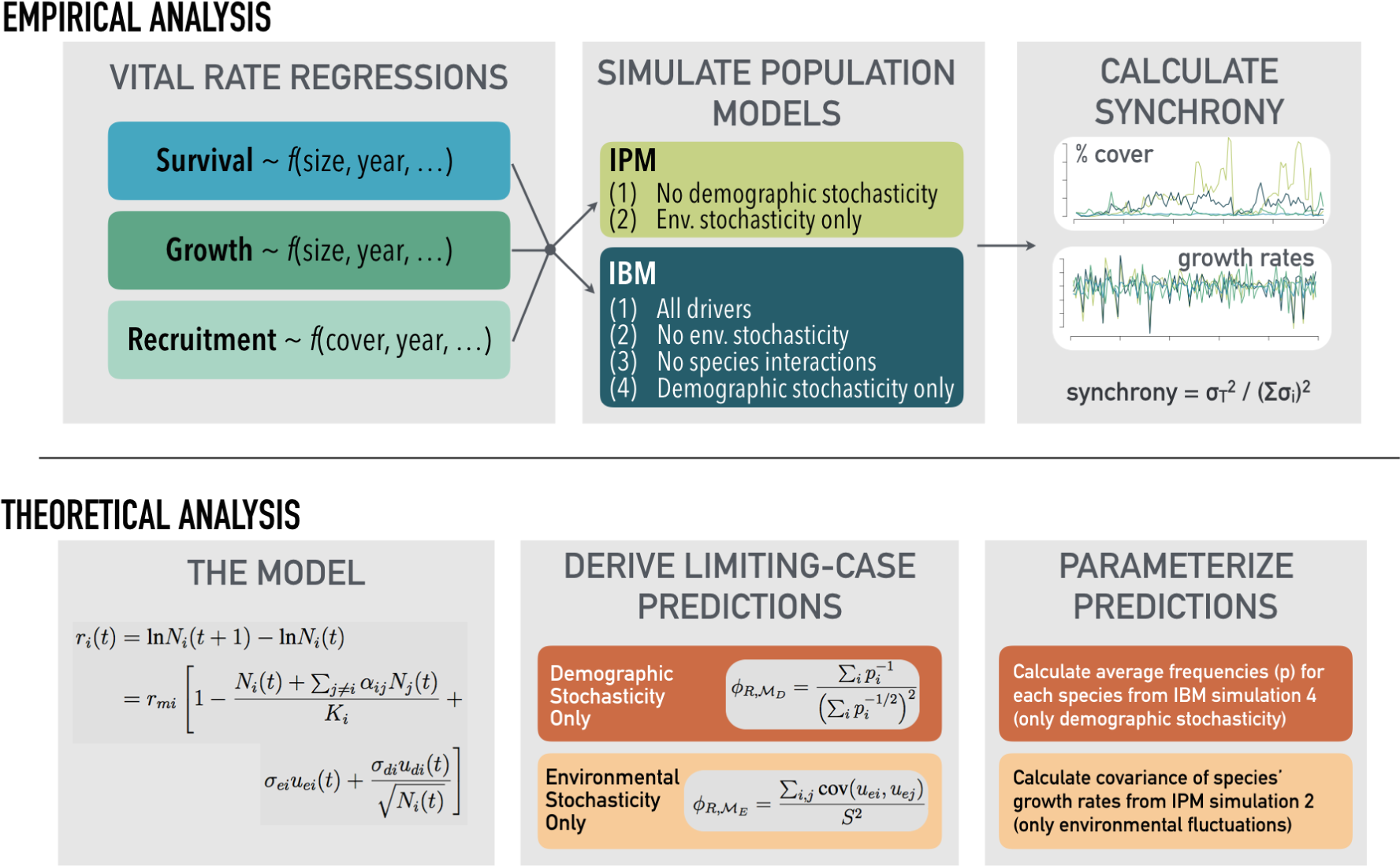
Diagram of our coupled theoretical-empirical approach. We followed this workflow for each of our five focal communities.

To achieve our objectives, we first refine theory that has been used to predict the effects of species richness on ecosystem stability (de Mazancourt et al. 2013) and species synchrony (Loreau and de Mazancourt 2008) to generate predictions of community-wide species synchrony under two limiting cases. We then confront our theoretical predictions with simulations from the empirically-based population models. Second, we use the multi-species population models to perform simulation experiments that isolate the effects of environmental stochasticity, demographic stochasticity, and interspecific interactions on community-wide species synchrony. Given that our population models capture the essential features of community dynamics important to synchrony (density-dependence, and demographic and environmental stochasticity), and that these models successfully reproduce observed community dynamics (Chu and Adler 2015), perturbing the models can reveal the processes that determine synchrony of dominant species in our focal grassland communities.

## THEORETICAL MODEL

### The model

While existing theory has identified environmental responses, species interactions, and demographic stochasticity as the drivers of the temporal synchrony, we do not have a simple expression to predict synchrony in a particular community with all factors operating simultaneously. However, we can derive analytical predictions for species synchrony under special limiting cases. The limiting case predictions we derive serve as baselines to help us interpret results from empirically-based simulations (described below). We focus on synchrony of per capita growth rates, rather than abundances, because growth rates represent the instantaneous response of species to the environment and competition, and are less susceptible to the legacy effects of drift and disturbance (Loreau and de Mazancourt 2008). We present equivalent results for synchrony of species abundances in the Appendix S1, and show that they lead to the same overall conclusions as synchrony of per capita growth rates.

Following Loreau and de Mazancourt (2008) and de Mazancourt et al. (2013), we define population growth, ignoring observation error, as

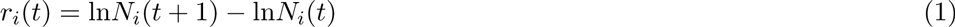

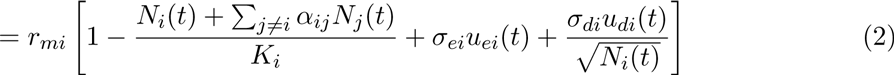

where *N_i_*(*t*) is the biomass of species *i* in year *t*, and *r_i_*(*t*) is its population growth rate in year *t*. *r_mi_* is species *i*’s intrinsic rate of increase, *K_i_* is its carrying capacity, and *α_ij_* is the interspecific competition coefficient representing the effect of species *j* on species *i*. Environmental stochasticity is incorporated as *σ_ei_u_ei_*(*t*), where 
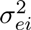
 is the temporal variance of species *i*’s response to the environment and *u_ei_*(*t*) is a normal variable with unit variance that is independent among species but may be correlated. The product, *σ_ei_u_ei_*(*t*), is species *i*’s environmental response. Demographic stochasticity arises from variations in births and deaths among individuals (e.g., same states, different fates), and is included in the model as a first-order, normal approximation (Lande et al. 2003, de Mazancourt et al. 2013). 
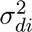
 is the demographic variance (i.e., the intrinsic demographic stochasticity of species *i*) and *u_di_*(*t*) are independent normal variables with zero mean and unit variance that allow demographic stochasiticity to vary through time.

To derive analytical predictions we solved a first-order approximation of Equation 2 (de Mazancourt et al. 2013 and Appendix S1). Due to the linear approximation approach, our analytical predictions will likely fail in communities where species exhibit large fluctuations due to limit cycles and chaos (Loreau and de Mazancourt 2008). Indeed, one of the advantages of focusing on growth rates rather than abundances is that growth rates are more likely to be well-regulated around an equilibrium value, if the long-term average of a species’ growth rate is relatively small (e.g., *r* < 2).

### Predictions

Our first prediction assumes no interspecific interactions, no environmental stochasticity, identical intrinsic growth rates, and that demographic stochasticity is operating but all species have identical demographic variances. This limiting case, 
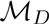
, represents a community where dynamics are driven by demographic stochasticity alone. Our prediction for the synchrony of per capita growth rates for 
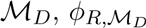
, is

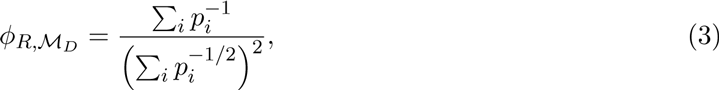

where *p_i_* is the average frequency of species *i*, *p_i_* = *N_i_*/*N_T_*. When all species have identical abundances and *p_i_* = 1/*S*, where *S* is species richness, synchrony equal 1/*S* (Loreau and de Mazancourt 2008).

Our second limiting case assumes only environmental stochasticity is operating 
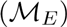
. Thus, we assume there are no interspecific interactions, demographic stochasticity is absent, intrinsic growth rates are identical, and environmental variance is identical for all species. Our prediction for the synchrony of per capita growth rates for 
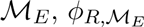
, is

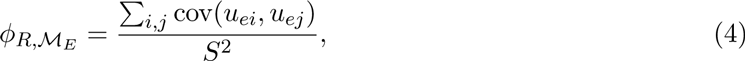

where cov(*u_ei_*, *u_ej_*) is the standardized covariance of environmental responses between species *i* and species *j*.

Confronting our theoretical predictions with data requires estimates of species dynamics of large populations (no demographic stochasticity) growing in isolation (no interspecific interactions) to calculate the covariance of species’ environmental responses. To estimate environmental responses in natural communities, we turn to our population models built using long-term demographic data.

## EMPIRICAL ANALYSIS

### Materials and methods

#### Data

We use long-term demographic data from five semiarid grasslands in the western United States (described in detail by Chu and Adler 2015). Each site includes a set of 1-m^2^ permanent quadrats within which all individual plants were identified and mapped annually using a pantograph (Hill 1920). The resulting mapped polygons represent basal cover for grasses and canopy cover for shrubs. Data come from the Sonoran desert in Arizona (Anderson et al. 2012), sagebrush steppe in Idaho (Zachmann et al. 2010), southern mixed prairie in Kansas (Adler et al. 2007), northern mixed prairie in Montana (Anderson et al. 2011), and Chihuahuan desert in New Mexico (Anderson et al. in preparation, Chu and Adler 2015) (Table 1).

**Table 1:**
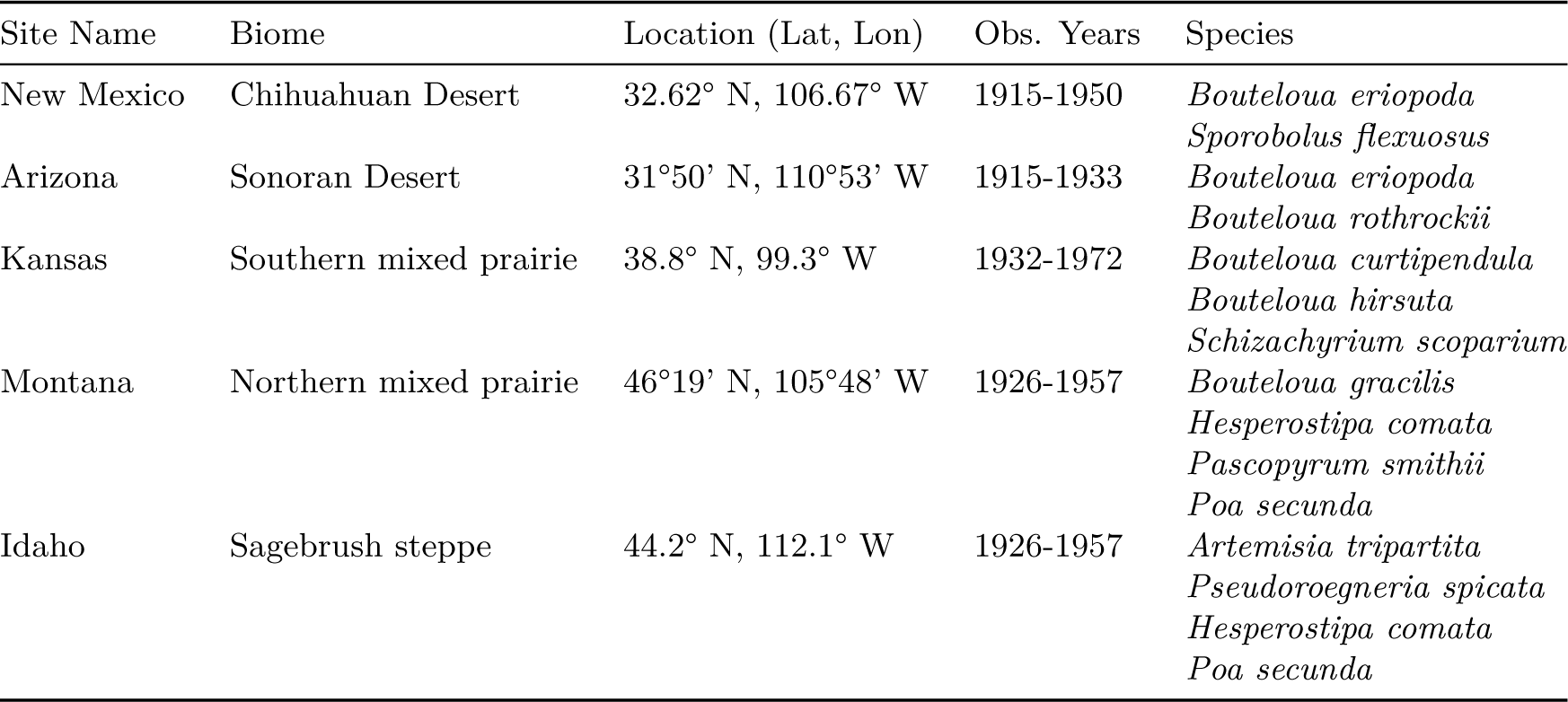
Site descriptions and focal species.

#### Calculating observed synchrony

The data consist of records for individual plant size in quadrats for each year. To obtain estimates of percent cover for each focal species in each year, we summed the individual-level data within quadrats and then averaged percent cover, by species, over all quadrats. We calculated per capita growth rates as log(*x_t_*) − log(*x*_*t*−1_), where *x* is species’ percent cover in year *t*. Using the community time series of per capita growth rates or percent cover, we calculated community synchrony using the metric of Loreau and de Mazancourt (2008) in the ‘synchrony’ package (Gouhier and Guichard 2014) in R (R Core Team 2013). Specifically, we calculated synchrony as

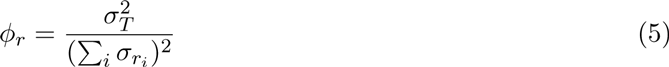

where 
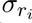
 is the temporal standard deviation of species *i’*s per capita population growth rate (*r_i_*) and 
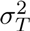
 is the temporal variance of the aggregate community-level growth rate. *ϕ* ranges from 0 at perfect asynchrony to 1 at perfect synchrony (Loreau and de Mazancourt 2008). We use the same equation to calculate observed synchrony of species’ percent cover, which we present to relate our results to previous findings, even though we focus on synchrony of growth rates in our model simulations (see below).

#### Fitting statistical models

Vital rate regressions are the building blocks of our dynamic models: an integral projection model (IPM) and an individual-based model (IBM). We followed the approach of Chu and Adler (2015) to fit statistical models for survival, growth, and recruitment (see Appendix S1 for full details). We modeled survival probability of each genet as function of genet size, temporal variation among years, permanent spatial variation among groups of quadrats, and local neighborhood crowding from conspecific and heterospecific genets. Regression coefficients for the effect of crowding by each species can be considered a matrix of interaction coefficients whose diagonals represent intraspecific interactions and whose off-diagonals represent interspecific interactions (Adler et al. 2010). These interaction coefficients can take positive (facilitative) or negative (competitive) values. We modeled growth as the change in size of a genet from one year to the next, which depends on the same factors as the survival model. We fit the survival and growth regressions using INLA (Rue et al. 2014), a statistical package for fitting generalized linear mixed effects models via approximate Bayesian inference (Rue et al. 2009), in R (R Core Team 2013). Spatial (quadrat groupings) variation was treated as a random effect on the intercept and temporal (interannual) variation was treated as random effects on the intercept and the effect of genet size in the previous year (Appendix S1).

Interspecific and intraspecific crowding, which represent species interactions, can be included as fixed effects or as random effects that vary each year. Strong year-to-year variation in these crowding effects would indicate a statistical interaction between environmental conditions and species interactions. We tested for such a dynamic by comparing models with and without random year effects on crowding. Based on the results, we decided to treat crowding as a fixed effect without a temporal component because most 95% credible intervals for random year effects on crowding broadly overlapped zero and, in a test case, including interannual variation in crowding did not change our results.

We modeled recruitment at the quadrat scale, rather than the individual scale, because the original data do not attribute new genets to specific parents (Chu and Adler 2015). Our recruitment model assumes that the number of recruits produced in each year follows a negative binomial distribution with the mean dependent on the cover of the parent species, permanent spatial variation among groups, temporal variation among years, and inter-and intraspecific interactions as a function of total species’ cover in the quadrat. We fit the recruitment model using a hierarchical Bayesian approach implemented in JAGS (Plummer 2003) using the ‘rjags’ package (Plummer 2014) in R (R Core Team 2013). Again, temporal and spatial variation were treated as random effects.

#### Building dynamic multi-species models

Once we have fit the vital rate statistical models, building the population models is straightforward. For the IBM, we initialize the model by randomly assigning plants spatial coordinates, sizes, and species identities until each species achieves a density representative of that observed in the data. We then project the model forward by using the survival regression to determine whether a genet lives or dies, the growth regression to calculate changes in genet size, and the recruitment regression to add new individuals that are distributed randomly in space. Crowding is directly calculated at each time step since each genet is spatially referenced. Environmental stochasticity is not an inherent feature of IBMs, but is easily included since we fit year-specific temporal random effects for each vital rate regression. To include temporal environmental variation, at each time step we randomly choose a set of estimated survival, growth, and recruitment parameters specific to one observation year. For all simulations, we ignore the spatial random effect associated with variation among quadrat groups, so our simulations represent an average quadrat for each site.

The IPM uses the same vital rate regressions as the IBM, but it is spatially implicit and does not include demographic stochasticity. Following Chu and Adler (2015), we use a mean field approximation that captures the essential features of spatial patterning to define the crowding index at each time step (Supporting Online Information). Temporal variation is included in exactly the same way as for the IBM. For full details on the IPMs we use, see Chu and Adler (2015).

#### Simulation experiments

We performed simulation experiments where drivers (environmental stochasticity, demographic stochasticity, or interspecific interactions) were removed one-by-one and in combination. To remove interspecific interactions, we set the off-diagonals of the interaction matrix for each vital rate regression to zero. This retains intraspecific interactions, and thus density-dependence, and results in simulations where species are growing in isolation. We cannot definitively rule out the effects of species interactions on all parameters, meaning that a true monoculture could behave differently than our simulations of a population growing without interspecific competitors. To remove the effect of a fluctuating environment, we removed the temporal (interannual) random effects from the regression equations. To remove the effect of demographic stochasticity, we use the IPM rather than the IBM because the IPM does not include demographic stochasticity (demographic stochasticity cannot be removed from the IBM). Since the effect of demographic stochasticity on population dynamics depends on population size (Lande et al. 2003), we can control the strength of demographic stochasticity by simulating the IBM on areas (e.g. plots) of different size. Results from an IBM with infinite population size would converge on results from the IPM. Given computational constraints, the largest landscape we simulate is a 25 m^2^ plot.

We conducted the following six simulation experiments: (1) IBM: All drivers (environmental stochasticity, demographic stochasticity, or interspecific interactions) present; (2) IPM: Demographic stochasticity removed; (3) IBM: Environmental stochasticity removed; (4) IBM: Interspecific interactions removed; (5) IPM: Interspecific interactions and demographic stochasticity removed; (6) IBM: Interspecific interactions and environmental stochasticity removed. We did not include a simulation with only interspecific interactions because our population models run to deterministic equilibriums in the absence of environmental or demographic stochasticity. We ran IPM simulations for 2,000 time steps, after an initial 500 iteration burn-in period. This allowed species to reach their stable size distribution. We then calculated yearly per capita growth rates from the simlated time series, and then caluclated the synchrony of species’ per capita growth rates over 100 randomly selected contiguous 50 time-step sections.

We ran IBM simulations for 100 time steps, and repeated the simulations 100 times for each simulation experiment. From those, we retained only the simulations in which no species went extinct due to demographic stochasticity. Synchrony of per capita growth rates was calculated over the 100 time steps for each no extinction run within a model experiment. In the IBM simulations, the strength of demographic stochasticity should weaken as population size increases, meaning that synchrony should be less influenced by demographic stochasticity in large populations compared to small populations. To explore this effect, we ran simulations (4) and (6) on plot sizes of 1, 4, 9, 16, and 25 m^2^. All other IBM simulations were run on a 25 m^2^ landscape to most closely match the implicit, large spatial scale of the IPM simulations.

Our simulations allow us to quantify the relative importance of environmental responses, species interactions, and demographic stochasticity by comparing the simulated values of community-wide species synchrony. The simulation experiments also allow us to test our theoretical predictions. First, in the absence of interspecific interactions and demographic stochasticity, populations can only fluctuate in response to the environment. Therefore, we can use results from simulation (5) to estimate the covariance of species’ responses to the environment (cov(*u_ie_*, *u_je_*)) and parameterize Equation 4. Parameterizing Equation 3 does not require simulation output because the only parameters are the species’ relative abundances. Second, simulations (5) and (6) represent the simulated version of our limiting case theoretical predictions. This approach to testing theoretical predictions may seem circular, but recall we derived the predictions using strict assumptions about equivalence in species’ growth rates, environmental response variances, and demographic variances. Our empirically-based models do not make these assumptions. Comparisons between parameterized predictions and simulated synchrony reveal whether the assumptions we must make to derive analytical predictions hold in the natural community our model represents.

### Results

Observed synchrony of species’ per capita growth rates at our study sites range from 0.36 to 0.89 and synchrony of percent cover ranged from 0.15 to 0.92 (Table 2). Synchrony of per capita growth rates and CV of percent cover are positively correlated (Pearson’s *ρ* = 0.72, *N* = 5). For all five communities, species synchrony from IPM and IBM simulations closely approximated observed synchrony (Fig. S1). IBM-simulated synchrony is consistently, but only slightly, lower than IPM-simulated synchrony (Fig. S1), likely due to the desynchronizing effect of demographic stochasticity.

**Table 2:** Observed synchrony among species’ per capita growth rates (*ϕ_R_*), observed synchrony among species’ percent cover (*ϕ_C_*), the coefficient of variation of total community cover, and species richness for each community. Species richness values reflect the number of species analyzed from the community, not the actual richness.

| Site | $\phi_R$ | $\phi_C$ | CV of Total Cover | Species richness |
| --- | --- | --- | --- | --- |
| New Mexico | 0.86 | 0.92 | 0.51 | 2 |
| Arizona | 0.89 | 0.80 | 0.47 | 2 |
| Kansas | 0.54 | 0.15 | 0.30 | 3 |
| Montana | 0.53 | 0.54 | 0.52 | 4 |
| Idaho | 0.36 | 0.18 | 0.19 | 4 |

Across the five communities, our limiting case predictions closely matched synchrony from the corresponding simulation experiment (Fig. 2 and Table S1). The correlation between our analytical predictions and simulated synchrony was 0.97 for 
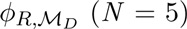
 and 0.997 for 
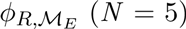
. The largest difference between predicted and simulated synchrony was 0.05 in New Mexico for 
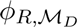
 (Table S1).

**Figure 2:**
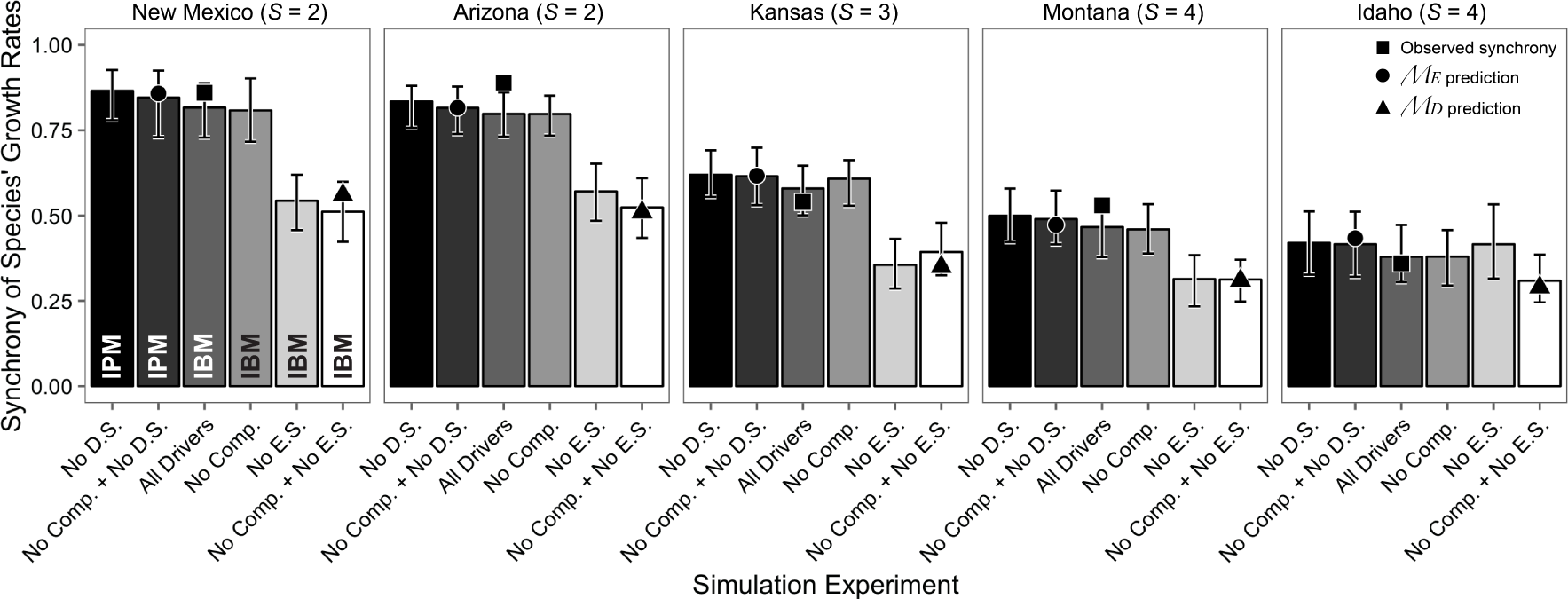
Community-wide species synchrony of per capita growth rates from model simulation experiments. Synchrony of species’ growth rates for each study area are from simulation experiments with demographic stochasticity, environmental stochasticity, and interspecific interactions present (“All Drivers”), demographic stochasticity removed (“No D.S.”), environmental stochasticity removed (“No E.S.”), interspecific interactions removed (“No Comp.”), interspecific interactions and demographic stochasticity removed (“No Comp. + No D.S.”), and interspecific interactions and environmental stochasticity removed (“No Comp. + No E.S.”). Abbreviations within the bars for the New Mexico site indicate whether the IBM or IPM was used for a particular simulation. Error bars represent the 2.5% and 97.5% quantiles from model simulations. All IBM simulations shown in this figure were run on a 25 m^2^ virtual landscape. Points show observed and predicted synchrony aligned with the model simulation that corresponds with each observation or analytical predicion.

Simulation experiments revealed that removing environmental fluctuations has the largest impact on synchrony, leading to a reduction in synchrony of species growth rates in four out of five communities (Fig. 2). Removing environmental fluctuations (“No E.S” simulations) decreased synchrony by 33% in Arizona, 48% in Kansas, 39% in Montana, and 40% in New Mexico. Only in Idaho did removing environmental fluctuations cause an increase in synchrony (Fig. 2), but the effect was small (9% increase; Table S2). Overall, species’ temporal random effects in the statistical vital rate models are positively, but not perfectly, correlated (Table S3). These temporal random effects represent environmental responses, meaning that positively correlated temporal random effects indicate positively correlated environmental responses.

Species interactions are weak in these communities (Table S4 and Chu and Adler 2015), and removing interspecific interactions had little effect on synchrony (Fig. 2; “No Comp.” simulations). Removing interspecific interactions caused, at most, a 5% change in synchrony (Fig. 2 and Table S2). Removing demographic stochasticity (“No D.S.” simulations) caused synchrony to increase slightly in all communities (Fig. 2), with an average 6% increase over synchrony from IBM simulations on a 25m^2^ area.

In IBM simulations, the desynchronizing effect of demographic stochasticity, which increases as population size decreases, modestly counteracted the synchronizing force of the environment, but not enough to lower synchrony to the level observed when only demographic stochasticity is operating (Fig. 3). In the largest, 25 m^2^ plots, synchrony was driven by environmental stochasticity (e.g., 
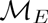
). At 1 m^2^, synchrony reflected the combined effects of demographic stochasticity and environmental stochasticity (e.g., simulated synhcrony fell between 
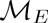
 and 
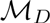
). For context, population sizes increased from an average of 17 individuals per community in 1 m to an average of 357 individuals per community in 25 m^2^ IBM simulations.

**Figure 3:**
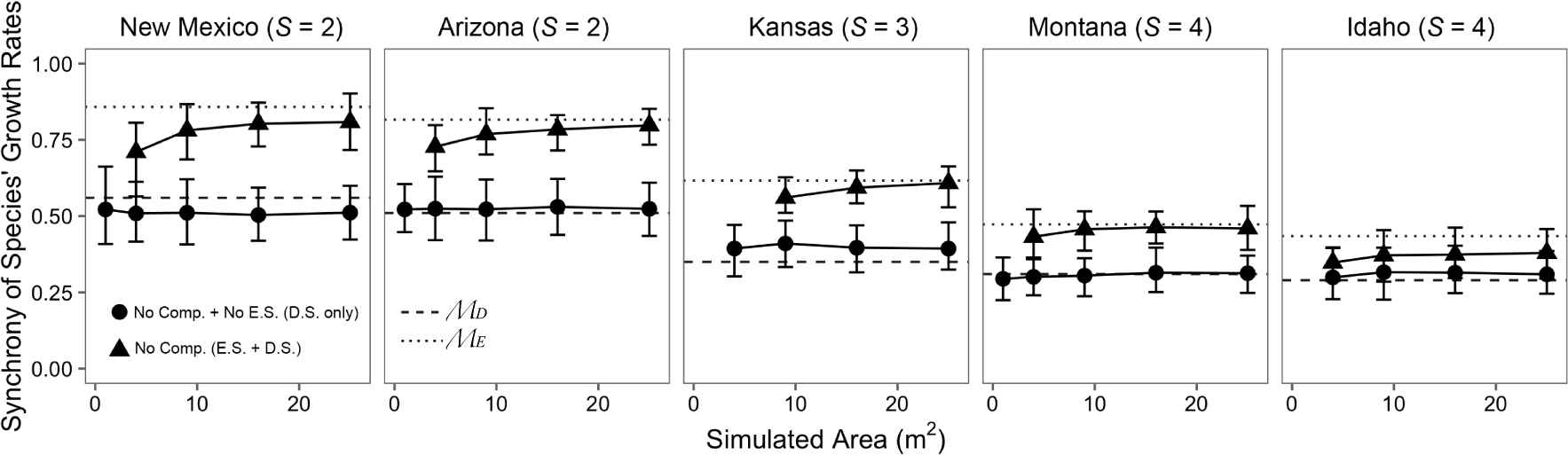
Synchrony of species’ growth rates for each study area from IBM simulations across different landscape sizes when only demographic stochasticity is present (“No Comp. + No E.S. (D.S. Only)”) and when environmental stochasticity is also present (“No Comp. (D.S. + E.S.)”). The horizontal lines show the analytical predictions 
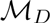
 (dashed line) and 
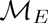
 (dotted line). The strength of demographic stochasticity decreases as landscape size increases because population sizes also increase. Theoretically, “No Comp. + No E.S. (D.S. Only)” simulations should remain constant across landscape size, whereas “No Comp. (D.S. + E.S.)” simulations should shift from the 
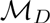
 prediction to the 
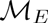
 prediction as landscape size, and thus population size, increases, but only if demographic stochasticity it strong enough to counteract environmental forcing. Error bars represent the 2.5% and 97.5% quantiles from model simulations.

Results for synchrony of percent cover are qualitatively similar, but more variable and less consistent with analytical predictions and observed synchrony (Appendix S1, Figs. S2-S3).

## DISCUSSION

Our study produced four main findings that were generally consistent across five natural plant communities: (1) limiting-case predictions from the theoretical model were well-supported by simulations from the empirical models; (2) demographic stochasticity decreased community synchrony, as expected by theory, and its effect was largest in small populations; (3) environmental fluctuations increased community synchrony relative to simulations in constant environments because species-specific responses to the environment were positively, though not perfectly, correlated; and (4) interspecific interactions were weak and therefore had little impact on community synchrony. We also found that analyses based on synchrony of species’ percent cover, rather than growth rates, were uninformative (Figs. S2-S3) since the linear approximation required for analytical predictions is a stronger assumption for abundance than growth rates, especially given relatively short time-series (Appendix S1). Thus, our results provide further evidence that it is difficult to decipher mechanisms of species synchrony from abundance time series, as expected by theory (Loreau and de Mazancourt 2008). Observed synchrony of per capita growth rates was positively correlated with the variability of percent cover across our focal communities, which confirms that we are investigating an important process underlying ecosystem stability.

### Simulations support theoretical predictions

Our theoretical predictions were derived from a simple model of population dynamics and required several restrictive assumptions, raising questions about their relevance to natural communities. For example, the species in our communities do not have equivalent environmental and demographic variances (Figs. S4-S7), as required by our predictions. However, the theoretical predictions closely matched results from simulations of population models fit to long-term data from natural plant communities (Table S1). Strong agreement between our analytical predictions and the simulation results should inspire confidence in the ability of simple models to inform our understanding of species synchrony even in complex natural communities, and allows us to place our simulation results within the context of contemporary theory. However, whether the theoretical model adequately represents more complex communities remains unknown because our analysis was restricted to dominant species.

### Demographic stochasticity decreases synchrony in small populations

In large populations, removing demographic stochasticity had no effect on species synchrony (Fig. 2). In small populations, demographic stochasticity partially counteracted the synchronizing effects of environmental fluctuations and interspecific interactions on per capita growth rates, in agreement with theory (Loreau and de Mazancourt 2008). This is shown in Fig. 3, where IBM simulations with environmental forcing and demographic stochasticity have higher synchrony than simulations with only demographic stochasticity. The differences between simulations are smaller at lower population sizes because as the number of individuals decreases with area, the strength of demographic stochasticity increases, reducing the relative effect of environmental forcing. Even in small populations (e.g., 1 m^2^ lanscapes), however, demographic stochasticity was not strong enough to compensate the synchronizing effects of environmental fluctuations and match the analytical prediction where only demographic stochasticity is operating ( 
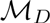
 in Fig. 3). These results confirm the theoretical argument of Loreau and de Mazancourt (2008) that independent fluctuations among interacting species in a non-constant environment should be rare. Only in the Idaho community does synchrony of per capita growth rates approach 
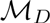
 in a non-constant environment (Fig. 3). This is most likely due to the strong effect of demographic stochasticity on the shrub *Artemisia tripartita* since even a 25 m^2^ quadrat would only contain a few individuals of that species.

Our analysis of how demographic stochasticity affects synchrony demonstrates that synchrony depends on the observation area. As the observation area increases, population size increases and the desynchronizing effect of demographic stochasticity lessens (Fig. 3). Thus, our results suggest that community-wide species synchrony will increase as the observation area increases, rising from 
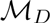
 to 
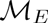
. Such a conclusion assumes, however, that species richness remains constant as observation area increases, which is unlikely (Taylor 1961). Recent theoretical work has begun to explore the linkage between ecosystem stability and spatial scale (Wang and Loreau 2014, 2016), and our results suggest that including demographic stochasticity in theoretical models of metacommunity dynamics may be important for understanding the role of species synchrony in determining ecosystem stability across spatial scales.

### Environmental fluctuations drive community synchrony

In large populations where interspecific interactions are weak, synchrony is expected to be driven exclusively by environmental fluctuations (Equation 4). Under such conditions community synchrony should approximately equal the synchrony of species’ responses to the environment (Loreau and de Mazancourt 2008). Two lines of evidence lead us to conclude that environmental fluctuations drive species synchrony in our focal plant communities. First, in our simulation experiments, removing interspecific interactions resulted in no discernible change in community-wide species synchrony of per capita growth rates (Fig. 2). Second, removing environmental fluctuations from simulations consistently reduced synchrony (Fig. 2). Our results lead us to conclude that environmental fluctuations, not species interactions, are the primary driver of community-wide species synchrony among the dominant species we studied. Given accumulating evidence that niche differences in natural communities are large (reviewed in Chu and Adler 2015), and thus species interactions are likely to be weak, our results may be general in natural plant communities.

In the Idaho community, removing environmental fluctuations did not cause a large decrease in synchrony. However, that result appears to be an artifact. Removing environmental variation results in a negative invasion growth rate for *A. tripartita*. Although we only analyzed IBM runs in which *A. tripartita* had not yet gone extinct, it was at much lower abundance than in the other simulation runs. When we removed *A. tripartita* from all simulations, the Idaho results conformed with results from all other sites: removing environmental stochasticity caused a significant reduction in species synchrony (Fig. S8). Our main results for Idaho (Fig. 2), with *A. tripartita* included, demonstrate how the processes that determine species synchrony interact in complex ways. *A. tripartita* has a facilitative effect on each grass species across all vital rates, except for a small competitive effect on *H. comata*’s survival probability (Tables S8-S10). At the same time, all the perennial grasses have negative effects on each other for each vital rate (Tables S8-S10). We know synchrony is affected by interspecific competition (Loreau and de Mazancourt 2008), but how facilitative effects manifest themselves is unknown. The interaction of facilitation and competition is clearly capable of having a large effect on species synchrony, and future theoretical efforts should aim to include a wider range of species interactions.

Environmental responses synchronized dynamics relative to a null expectation of independent species interactions (e.g., “No Comp. + No E.S.” simulations in Fig. 2), but observed and simulated synchrony was still less than one in all cases (Fig. 2). Synchrony was far from complete because of differences in species’ responses to interannual environmental variation. Many studies of ecosystem stability in semiarid grasslands focus on trad-offs among dominant functional types (Bai et al. 2004, Sasaki and Lauenroth 2011). Such groupings are based on the idea that ecologically-similar species will have similar responses to environmental fluctuations. At first glance our results may appear to support the grouping of perennial grasses in one functional type because their environmental responses were positively correlated. However, even though environmental responses among the dominant species we studied were similar, they were dissimilar enough to cause synchrony to be less than perfect (Fig. 2). The subtle differences among ecologically-similar dominant species do impact species synchrony and, ultimately, ecosystem stability. Ignoring such differences could mask important dynamics that underpin ecosystem functioning.

### Interspecific interactions had little impact on community synchrony

We expected community synchrony of per capita growth rates to decrease when we removed interspecific interactions (Loreau and de Mazancourt 2008). We found that community synchrony was virtually indistinguishable between simulations with and without interspecific interactions (Fig. 2). The lack of an effect of interspecific interactions on synchrony is in contrast to a large body of theoretical work that predicts a strong role for competition in creating compensatory dynamics (Tilman 1988) and a recent empirical analysis (Gross et al. 2014).

Our results do not contradict the idea that competition can lead to compensatory dynamics, but they do highlight the fact that interspecific competition must be relatively strong to influence species synchrony. The communities we analyzed are composed of species with very little niche overlap (Chu and Adler 2015) and weak interspecific interactions (Tables S1 and S3-S17). Mechanistic consumer-resource models (Lehman and Tilman 2000) and phenomenological Lotka-Volterra models (Lehman and Tilman 2000, Loreau and de Mazancourt 2013) both confirm that the effect of competition on species synchrony diminishes as niche overlap decreases. In that sense, our results are not surprising: interspecific interactions are weak, so of course removing them does not affect synchrony.

Our conclusion that species interactions have little impact on synchrony only applies to single trophic level communities. Species interactions almost certainly play a strong role in multi-trophic communities where factors such as resource overlap (Vasseur and Fox 2007), dispersal (Gouhier et al. 2010), and the strength of top-down control (Bauer et al. 2014) are all likely to affect community synchrony.

## CONCLUSIONS

Species-specific responses to temporally fluctuating environmental conditions is an important mechanism underlying asynchronous population dynamics and, in turn, ecosystem stability (Loreau and de Mazancourt 2013). When we removed environmental variation, we found that synchrony decreased in four out of the five grassland communities we studied (Fig. 2). A tempting conclusion is that our study confirms that compensatory dynamics are rare in natural communities, and that ecologically-similar species will exhibit synchronous dynamics (e.g., Houlahan et al. 2007). Such a conclusion misses an important subtlety. The perennial grasses we studied do have similar responses to the environment (Table S2), which will tend to synchronize dynamics. However, if community-wide synchrony among dominant species is less than perfect, as it is in all our focal communities, some degree of compensatory dynamics must be present (Loreau and de Mazancourt 2008). Even ecologically-similar species, which are sometimes aggregated into functional groups, have environmental responses that are dissimilar enough to limit synchrony. Subtle differences among dominant species ultimately determine ecosystem stability and should not be ignored.

## ACKNOWLEDGEMENTS

This work was funded by the National Science Foundation through a Postdoctoral Research Fellowship in Biology to ATT (DBI-1400370) and a CAREER award to PBA (DEB-1054040). CdM and ML were supported by the TULIP Laboratory of Excellence (ANR-10-LABX-41). We thank the original mappers of the permanent quadrats at each site and the digitizers in the Adler lab, without whom this work would not have been possible. Compute, storage, and other resources from the Division of Research Computing in the Office of Research and Graduate Studies at Utah State University are gratefully acknowledged. We also thank Patrick Venail and several anonymous reviewers who provided critical feedback that improved the manuscript.

